# HistoClean: Open-source Software for Histological Image Pre-processing and Augmentation to Improve Development of Robust Convolutional Neural Networks

**DOI:** 10.1101/2021.06.07.447339

**Authors:** Kris D. McCombe, Stephanie G. Craig, Amélie Viratham Pulsawatdi, Javier I. Quezada-Marín, Matthew Hagan, Simon Rajendran, Matthew P. Humphries, Victoria Bingham, Manuel Salto-Tellez, Richard Gault, Jacqueline A. James

## Abstract

The growth of digital pathology over the past decade has opened new research pathways and insights in cancer prediction and prognosis. In particular, there has been a surge in deep learning and computer vision techniques to analyse digital images. Common practice in this area is to use image pre-processing and augmentation to prevent bias and overfitting, creating a more robust deep learning model. Herein we introduce HistoClean; user-friendly, graphical user interface that brings together multiple image processing modules into one easy to use toolkit. In this study, we utilise HistoClean to pre-process images for a simple convolutional neural network used to detect stromal maturity, improving the accuracy of the model at a tile, region of interest, and patient level. HistoClean is free and open-source and can be downloaded from the Github repository here: https://github.com/HistoCleanQUB/HistoClean.

## 1. Introduction

The growth of digital image analysis in clinical pathology and its subsequent case for use in clinical medicine has been supported by the conception of open-source digital image analysis (DIA) software [1–3]. Use of machine learning from predetermined features allows for the development of DIA algorithms within these software environments. This allows bio-image analysts and consultant histopathologists to answer difficult, specific research questions in human tissue [4]. The subsequent introduction of deep learning has revolutionised the development of DIA algorithms [5]. This has enabled potential solutions to tumour and biomarker detection, as well as tumour subtyping [6,7]. However, these solutions require domain-specific knowledge relating to the deep learning methodology, as well as the awareness of hardware acceleration [8].

Consequently, open-source software to facilitate bio-image analysts without a background in computer vision to develop deep learning models have evolved [9,10]. Deep learning methodologies learn feature representations from the data without requiring predefined feature extraction. The resultant models can therefore be significantly more sensitive to dataset specific attributes, such as irregularities in staining, batch effects and the quality of the digital slide [11,12]. Use of image pre-processing and augmentation prior to developing deep learning models can regularise the input images, thereby, mitigating the potential for bias in the training of the CNN, or other deep learning models, and its independent validation [13–16]. Among these, the most common techniques include class-balancing [17], image normalisation [18], and image augmentation [19]. These techniques often involve the use of multiple coding libraries, which in turn requires knowledge of the documentation before implementation.

Herein we present HistoClean; an open-source, high-level, graphical user interface (GUI) for image pre-processing. HistoClean aims to complement other open-source software and deep-learning frameworks in the bio-image analysis ecosystem [18, 19]. HistoClean’s image pre-processing toolkit is divided into five functional modules based on computational methods frequently used in histological image pre-processing; image patching, whitespace thresholding, dataset balancing, image normalisation and image augmentation (Figure 1). These modules can be used independently or in combination with each other as the user requires. HistoClean brings together image pre-processing techniques from across multiple Python libraries. This simplifies the image preparation phase of deep-learning analysis in a way that is transparent and maintains data integrity.

**Figure 1.**
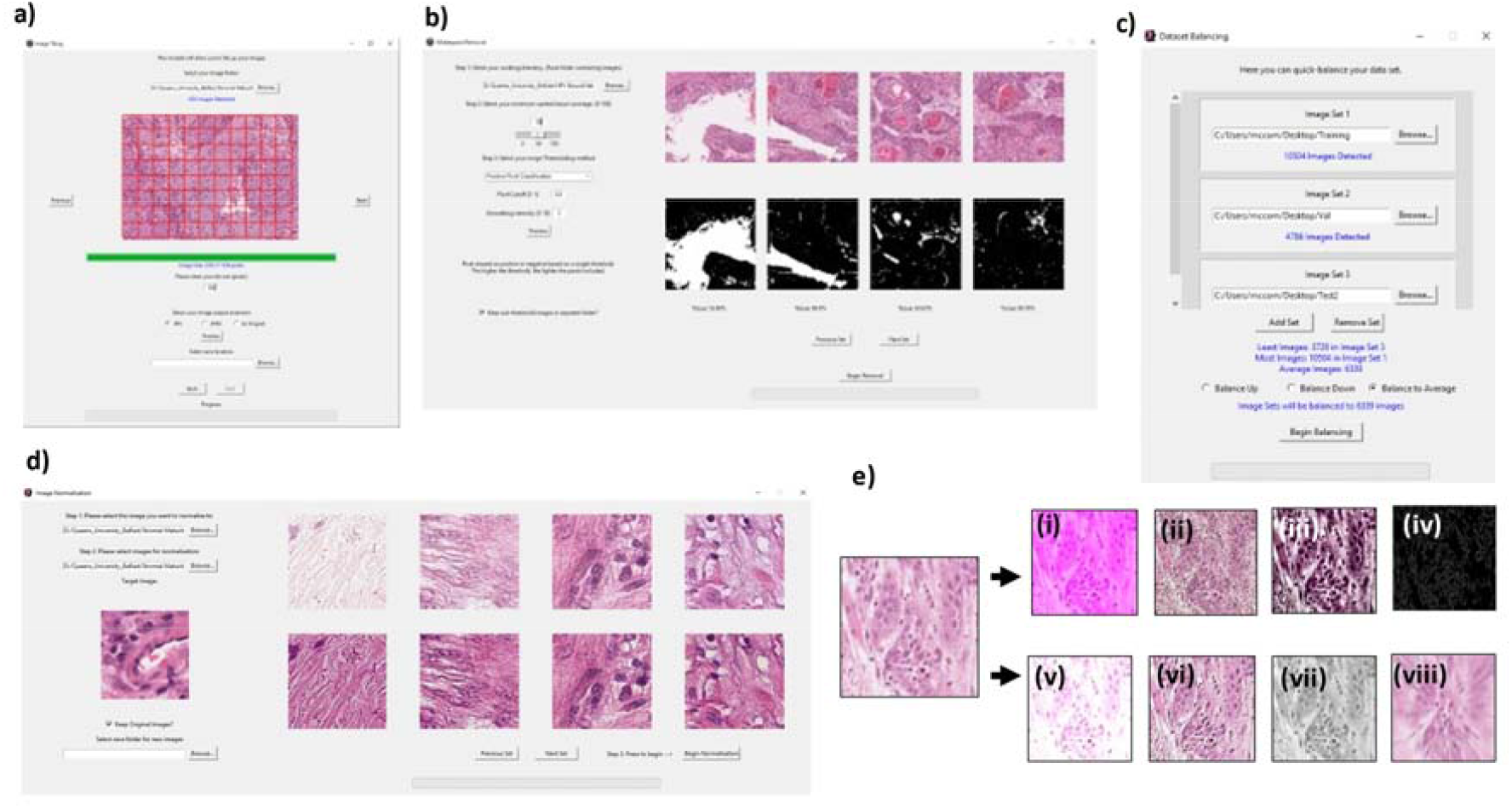
HistoClean, an all-in-one toolkit for the pre-processing of images for use in deep learning. Modules include (a) tools for generation of image tiles from larger images, which are executed within the image patching module (b), whitespace estimation and filtering, implemented in the white space removal module (d) image normalisation, which standardises the colour grading of the images and image augmentation techniques, which are implemented in the dataset balancing, image normalisation and image augmentation modules as demonstrated in the representative images shown (e). Examples of augmentation include but are not limited to: (i) Increased red value, (ii) Pixel Dropout (iii) Increased contrast (iv) Canny edge detection. (v) Brightness (vi) Embossing (vii) Greyscale (viii) Motion blur.

In this study, a practical example of how HistoClean can optimise input images for training CNNs to predict stromal maturity is described (Figure 2). In evaluating these models, we demonstrate the benefit of image pre-processing for deep learning, and introduce HistoClean as an open-source software solution to quickly implement and review these techniques.

**Figure 2.**
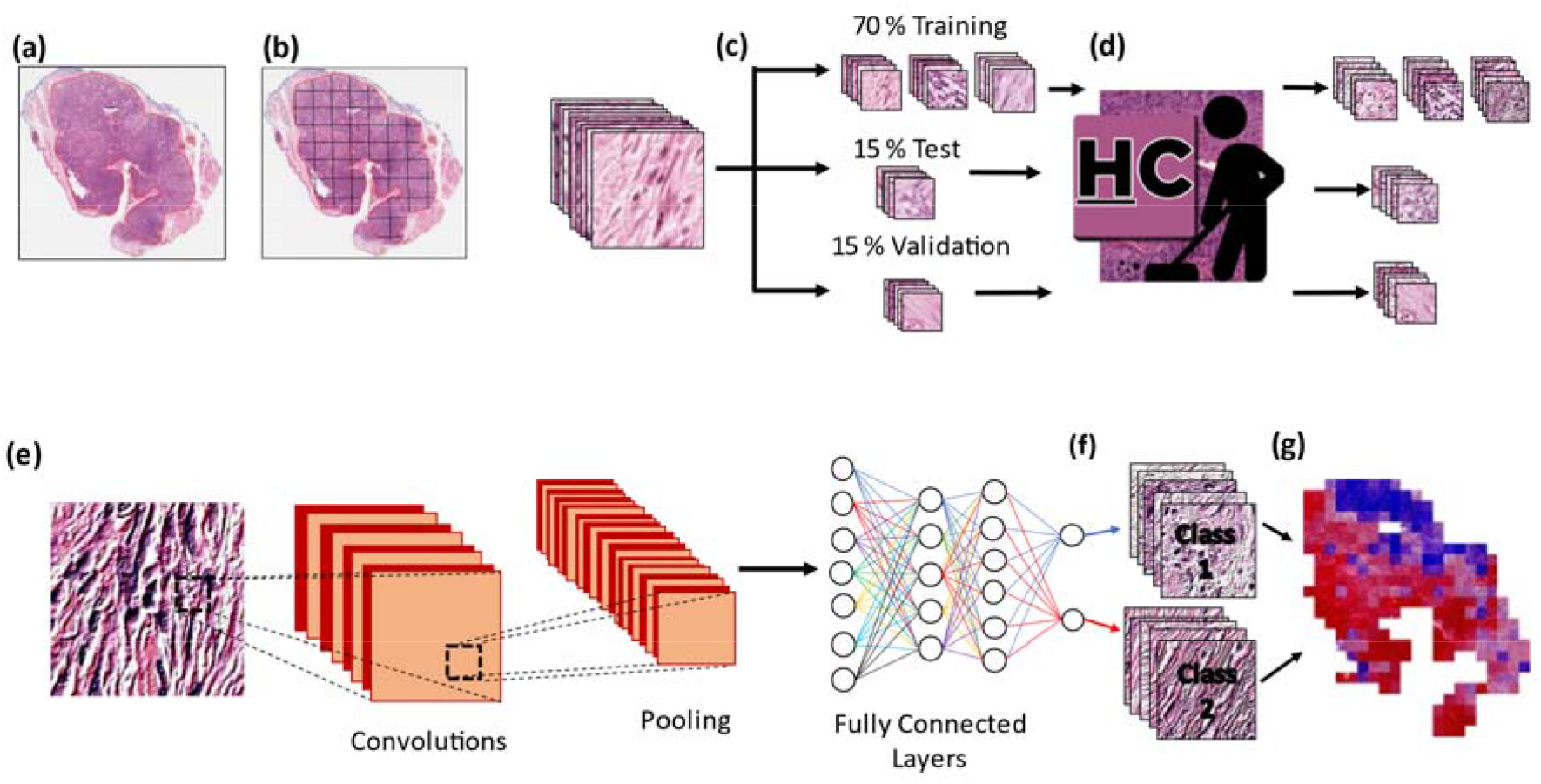
Use of HistoClean in the development of histology based convolutional neural networks. Slides are scanned at high-resolution, normally 200-400x magnification and are virtually annotated (as outlined in red) by a pathologist on a digital platform (a). Tiles of equal size are extracted from the virtual annotations (b). These tiles are independently sorted into training, test and validation datasets at a patient level (c). Image pre-processing and augmentation is conducted on the tiles using HistoClean where appropriate in the training, test and validation datasets in order to prepare tiles for use in a convolutional neural network (d). Within a typical convolutional neural network, each tile is fed through a series of convolutional and pooling layers in order to create feature maps to differentiate between the two classes (e). These feature maps are then fed through several fully connected layers which determine which class the images belong to (f). Each tile is assigned a value used for class prediction; the prediction values for each tile are then aggregated in order to provide a overall class prediction per patient (g).

The main contribution of this paper is the development of a novel, easy to use application for the rapid pre-processing and augmentation of image datasets for use in deep learning, image analysis pipelines.

## 2. Materials and Methods

### 2.1 HistoClean Application Development

HistoClean was developed using Anaconda3 and Python 3.8. Code was written using the PyCharm integrated developer environment. The GUI was developed using the Tkinter toolbox (v8.6). Initial development and testing of the software was performed on an Octane V laptop with an Intel Core i7-9700F 3.0GHz processor and 32GB Corsair 2400MHz SODIMM DDR4 RAM, with a Windows 10 operating system. The application was converted to a .exe program using the Pyinstaller Python package [21]. All testing was performed in the Windows 10 operating system. For ease of use it is recommended that images should be organised within directories corresponding to each image class. The application runs all processes on the CPU. No GPU is required. The application makes prominent use of multithreading, which scales to the number of cores in the CPU.

#### 2.1.1 Image Patching Module

CNN’s require input image tiles to have consistent dimensions [22]. For this reason, HistoClean includes an image patching module that utilises the Python library *Patchify* [23]. This module interface allows the user to create image tile subsets from a larger input image to their specification and provides real-time feedback of the output to the user, facilitating straightforward evaluation and adjustment (Figure 1a). This module can be used for block processing of *n* images organised within a common file directory. The user can select an output destination wherein the directory structure and naming conventions of the original images will be retained and populated with the requested image patches. The file names of these new image tiles are suffixed with their patch co-ordinates from the original image for reproducibility. Maintaining transparency in the pre-processing stages ensures that results can ultimately be traced back to their source ensuring that HistoClean does not damage original source data or impede data integrity and reproducibility.

#### 2.1.2 Tissue Thresholding Module

Most pathology-orientated CNN’s are developed to address questions within the tissue, therefore, an excess of whitespace in the input images may impair model development [24]. In order to address this issue and improve the quality of input image tiles, HistoClean includes a tissue thresholding module that allows the user to remove image tiles from their dataset based on a minimum threshold of approximate tissue coverage. The method outlined in this paper uses binary thresholding to determine the percentage of positive pixels, representing tissue, and null pixels, representing whitespace (Figure 3). Tissue coverage and relative intensity of the staining can vary significantly depending on any number of predisposing factors. Therefore, HistoClean’s module interface allows the user, in real time, to explore different thresholds for dichotomising these pixels into tissue vs whitespace. In addition, adaptive thresholding is available for each image as well as Otsu binarization [25]. All of these thresholding options come courtesy of the OpenCV Python library [26]. These processes generate a binary mask for each image which the GUI presents alongside the original image for review. Users can view five images simultaneously. Upon approval of an arbitrary threshold, images are removed or relocated based on user preference.

**Figure 3.**
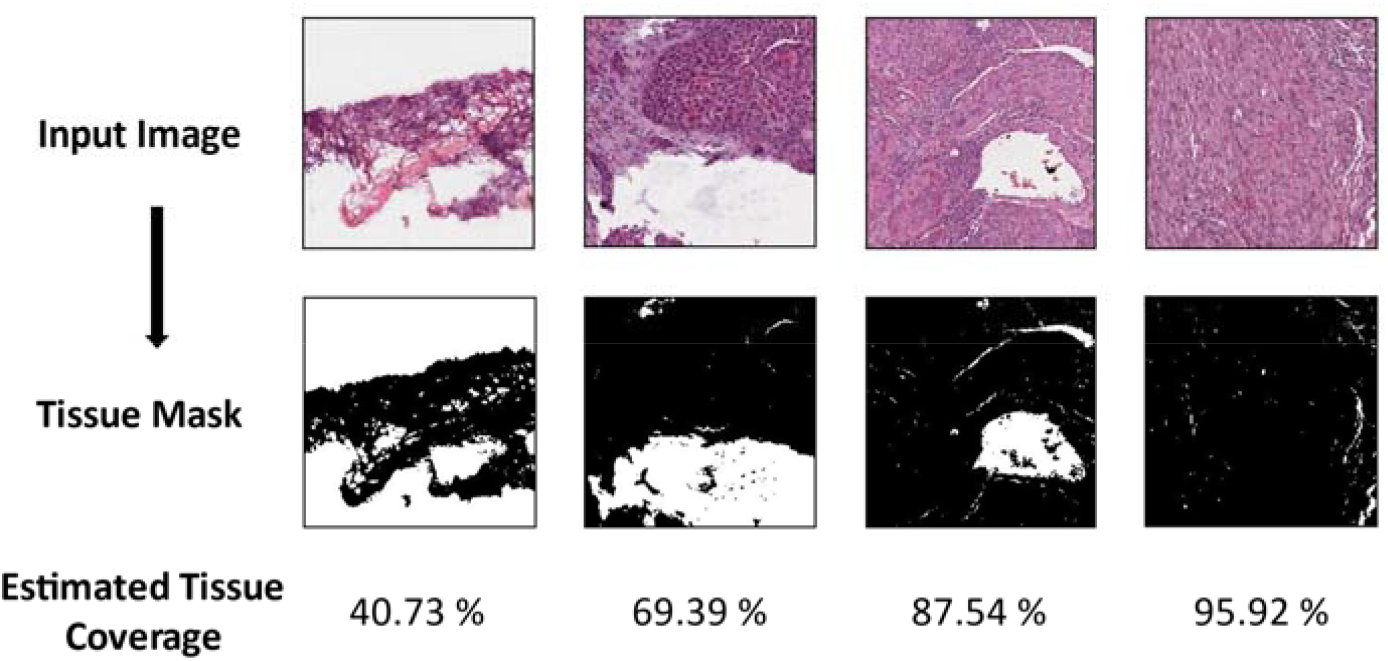
Four representative images demonstrating use of the positive pixel classifier method to estimate tissue coverage. All images were given the same cut-off (0.8). The top row contains the original images, with the bottom row showing the binary mask for tissue (black = tissue, white = whitespace). The bottom row shows the estimated tissue coverage within the image tile.

#### 2.1.3 HistoClean: class balancing module

Class balancing is essential to prevent class bias of data when developing deep learning models [27]. For this reason, HistoClean includes a class balancing module that enables the user to equalise the number of images per class prior to training of the CNN (Figure 1c). This requires that each class of images be provided in a separate directory by the user. The user can then decide to balance using three options: reducing the number of image tiles in each class to the smallest class, increasing the number of image tiles in each class based on the largest class, or balance the number of images in each class based on the average number of images in each class. The pre-requisite for using this functionality is that no class contains less than one eighth of the samples of the largest class. This pre-condition is reinforced through exception handling. This is to prevent duplicate images arising from repeated augmentations. If the user balances the samples through class reduction, the image tiles in the larger class-specific dataset are then relocated to a new directory, denoted as ‘Removed Images’, or are permanently deleted based on user preference. If class-size is balanced by the addition of image tiles, then a random assortment of image tiles equal to the difference between the largest class-specific image dataset are selected without replacement from within the smaller dataset(s). The random selections of image tiles are then augmented thus balancing the number of image tiles in that class by addition of ‘new’ image data. Image augmentation techniques are randomly selected from mirroring, clockwise rotation at 90°, 180° or 270°, or a combination of mirroring and a single rotation. This can create up to 7 unique images from a single image as required. A random number generator, seeded to the date and time of dataset balancing, determines the augmentation applied.

#### 2.1.4 Image Normalisation Module

Histological images possess unique image colour, contrasts, and brightness profiles. Batch effects in staining (Figure 4a) can significantly influence model performance [13]. Image normalisation can be used to bring uniformity to the images in the dataset by adjusting the range of pixel values of an input image, according to that of a target image [18]. For this reason, HistoClean includes an image normalisation module based on histogram matching from the Python library *scikit-image* [28]. Histogram matching works by comparing the cumulative histogram of pixel intensities from a target and an input image, before adjusting the pixel values of the input image according to the target image [29] (Figure 4b). HistoClean’s module interface allows the user to select a target image to normalise to and to review examples of the histogram-matched images before committing to image normalisation to *n* images organised within a folder. These are saved to a separate user-defined folder, or can replace the original images at the user’s discretion.

**Figure 4.**
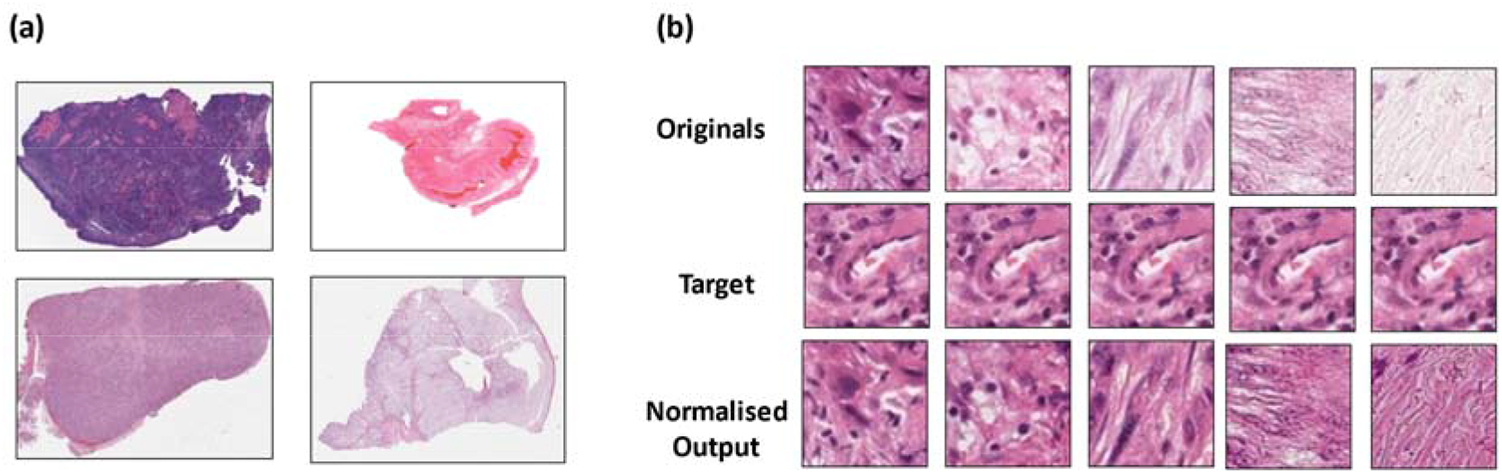
Image normalisation in histological images. Batch effects in haematoxylin and eosin staining and different staining protocols often leads to an inconsistent colour range in histological images as demonstrated by images taken from the TCGA head and neck diagnostic dataset (a). Demonstration of histogram normalisation to correct for the inconsistent colour range between samples while preserving histological architecture (b). The top row shows a selection of original un-normalised tiles, the middle row shows the target image and preferred colour range being normalised to and the bottom row shows the result of that normalisation.

#### 2.1.5 Image Augmentation and Pre-processing Module

It is not always possible to source large collections of histological images in the pursuit of developing deep learning models [30]. Image augmentation is a technique which that can be used for the artificial expansion of image datasets to provide more training examples. In addition, image pre-processing can be used to enhance features already present in an image dataset in order to provide more specific features for the CNN training [31]. By providing deep learning models with augmented data, the user can reduce the risk of overfitting and improve the generalisation ability of the CNN [30]. For this reason, HistoClean includes an image augmentation/pre-processing module based on the Python library *Imgaug* [32]. This allows the user to select, review and apply the most popular image augmentation techniques used in the development of CNNs to their image dataset in real-time (Figure 1e). These include adjusting the colour range, contrast, blur and sharpness, noise, pixel and channel dropout and more. Generated images files from augmentation are identifiable by their name, which incorporates the name of the root file from which the image derived so as to maintain data integrity.

### 2.2 Patient samples

Ethical approval and access to diagnostic H&E stained slides from a retrospective cohort of oropharyngeal squamous cell carcinomas (OPSCC) for stromal maturity prediction by artificial intelligence was granted via the Northern Ireland Biobank (OREC 16/NI/0030; NIB19/0312) [33]. Briefly, patients with a primary oropharyngeal cancer diagnosed between 2000-2011 were identified and their diagnostic H&E retrieved from the Belfast Health and Social Care Trust courtesy of the Northern Ireland Biobank. All slides were digitised using a Leica Aperio AT2 at 40x magnification (0.25μm / pixel). Virtual slides were saved in a .svs file format and imported into the open-source image analysis tool QuPath (v0.1.2) [1] to enable image annotation by a qualified histopathologist.

### 2.3 Classification of stromal maturity

Using DIA software QuPath (v0.1.2), a trained pathologist reviewed all the diagnostic H&E slides from each case before identifying and annotating ROIs for classification of stromal maturity on the slide that most represented malignant OPSCC. Stroma maturity was determined as being either mature or immature for each ROI by visual review. This was conducted by the pathologist, along with two other blinded independent assessors based on previously published criteria [34,35]. Classification of mature stroma was defined by the presence of fine, regular, elongated collagen fibres organised with approximately parallel orientation. Conversely, immature stroma was defined by disorganised, random orientation of collagen fibres with and without the presence of edema and myxoid-like degeneration. Representative images of mature and immature stroma were created and used as reference criteria for all assessors prior to classification (Figure 5).

**Figure 5.**
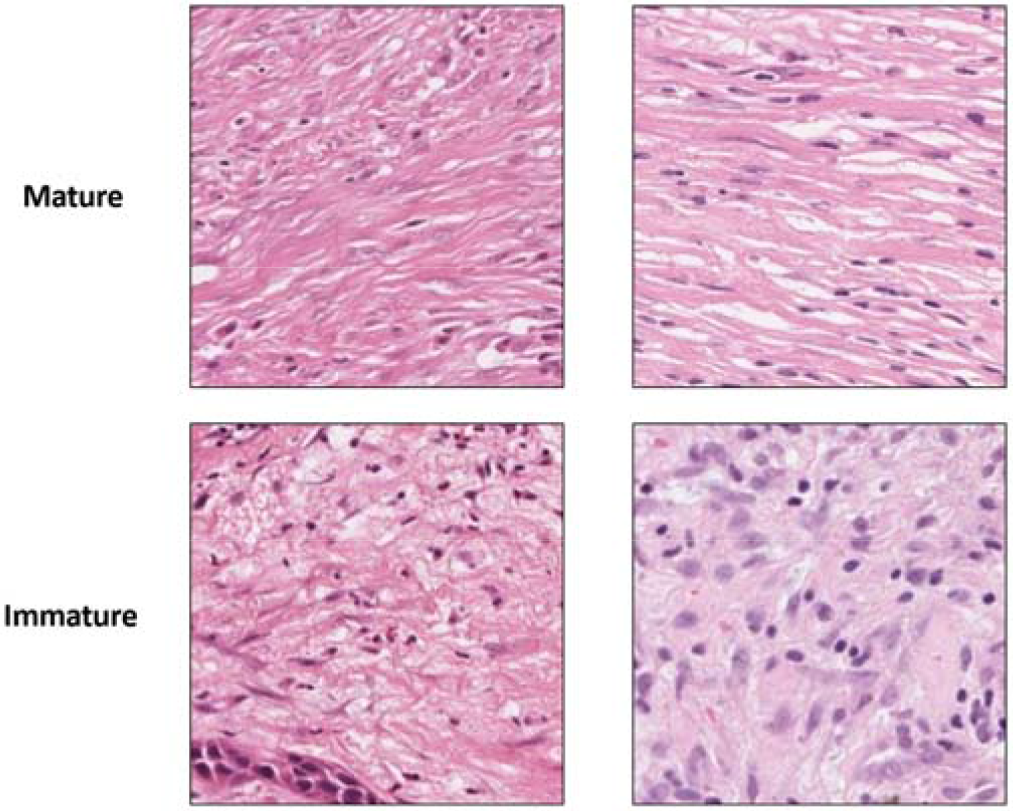
Reference images for mature (top row) and immature (bottom row) stroma randomly selected from the dataset. Images taken at 40x magnification and used as reference criteria during the manual classification of stromal maturity by the independent assessors in the study.

### 2.4 Image Set Preparation

Image tiles of size 250×250 pixels were extracted from the stromal regions of the annotated ROIs utilising QuPath’s scripting functions. Tiles were organised in separate directories for mature and immature stroma as determined by manual assessment. These were further grouped into directories representing each patient. Images were divided at a patient level into three sets. First, the training set, which consisted of 70% of the patients was used to train the CNNs. Second, the test set, which consisted of 15% of the patients was used to evaluate model performance during training. Lastly, the independent validation set consisting of the remaining 15% of patients. This did not influence the training of the model and was instead used to evaluate model performance. This produced the baseline “Unbalanced” image set. Images were organised in this way to account for intra-patient heterogeneity of stromal maturity. An entire heterogenous patient existed within the training, test or independent validation set and was not split among the three. This is to prevent the CNN from “recognising” patients between the three sets.

### 2.5 Image pre-processing using HistoClean

In order to demonstrate the benefit of image pre-processing for the development of robust CNN’s, seven independent image datasets were produced from the baseline image set. These utilised a combination of class balancing, image normalisation and pre-processing (Table 1). Class balancing augmented the smaller image class to provide the same number of images as the larger class. This option was chosen as reducing the larger class down, would have resulted in a lesser volume of images for training, harming model accuracy. Image pre-processing was limited to embossing of the images (Intensity = 2, Alpha = 1) (Figure 6). The same target image was used in all normalised sets. All image manipulation was conducted prior to input in the CNN. The processes for creating all these image sets were timed. Augmentations were applied across the training, test and independent validation sets, with the exception of balancing, which was done across training and test sets only.

**Table 1.** Summary table of Histoclean modules used in each dataset. Columns denoted with an “X” show which modules were used.

| Dataset | Balancing | Normalisation | Augmentation |
| --- | --- | --- | --- |
| Unbalanced |  |  |  |
| Unbalanced Normalised |  | X |  |
| Unbalanced Embossed |  |  | X |
| Unbalanced Normalised Embossed |  | X | X |
| Balanced | X |  |  |
| Balanced Normalised | X | X |  |
| Balanced Embossed | X |  | X |
| Balanced Normalised Embossed | X | X | X |

**Figure 6.**
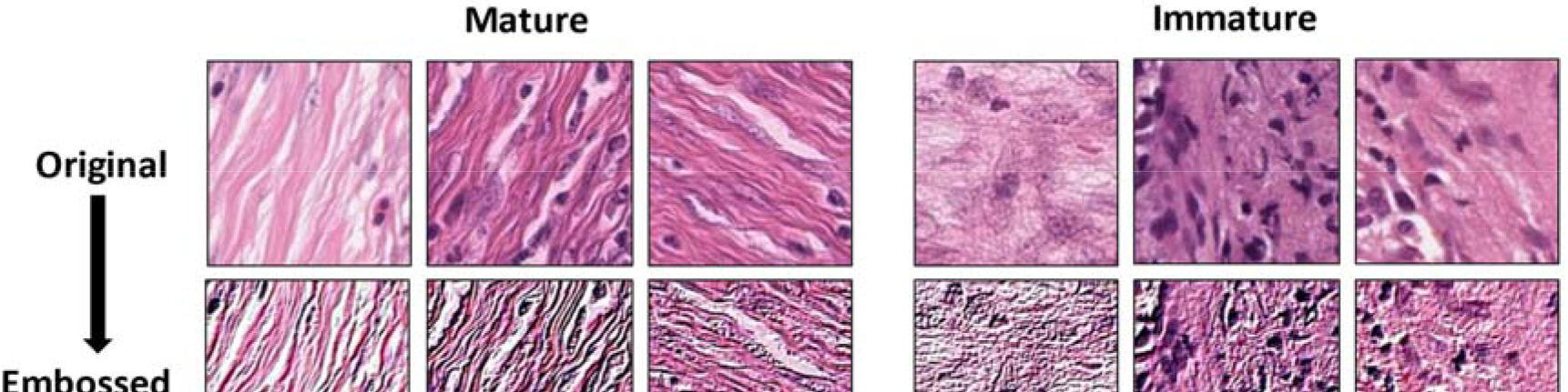
Demonstration of embossing on mature and immature tiles. The top row consists of the original images and the bottom row shows the effects of embossing. Embossing accentuates the difference between the two stroma maturities.

### 2.6 CNN Design

The CNNs used in these experiments were designed using PyTorch [36]. A core CNN architecture was established and trained independently on each of the 8 datasets from scratch. This network consists of five convolutional layers interlinked with five pooling layers (Figure 7). The output of the final pooling layer is then flattened and fed into two fully connected layers wherein stromal maturity is predicted using the softmax function in the final layer. Training was carried out for 200 epochs, with a batch size of 150. Adam Optimisation was used with a learning rate of 1e-6. Test batch size was set to 150 images. The outcome of the softmax function in the CNN produced a probability for each input image ranging from 0 (predicted mature) to 1 (predicted immature). Stromal maturity of the input images was classified as immature if the stromal maturity probability was greater or equal to 0.5, otherwise it was considered mature. After training on every fifth batch, the neural network calculated the accuracy and loss on a randomly selected test batch. If the test accuracy was greater than or equal to 65%, the weights and biases of the model were saved for further model evaluation. The weights and biases of the top 10 test batch accuracies were applied to the entire test set to get an improved evaluation of in-model performance. Only the model weights and biases that provided the top test accuracy were carried forward. These were then loaded to the CNN and applied to the independent validation image set. Stromal maturity probabilities at a ROI level were produced by majority voting of individual tile classifications. In patients with heterogeneous ROI classification of stromal maturity, majority voting of the ROIs was used to determine classification at a patient level. This was done to remain comparable with manual assessment. If the number of predicted stromal immature and mature ROI’s was equal the patient was considered to have mature stroma overall. To enable comparison of how different input images affected training of the CNN, batch size, learning rate, loss function and optimiser were all kept constant through all experiments. Full code for the CNN can be found at: (https://github.com/HistoCleanQUB/HistoClean)

**Figure 7.**
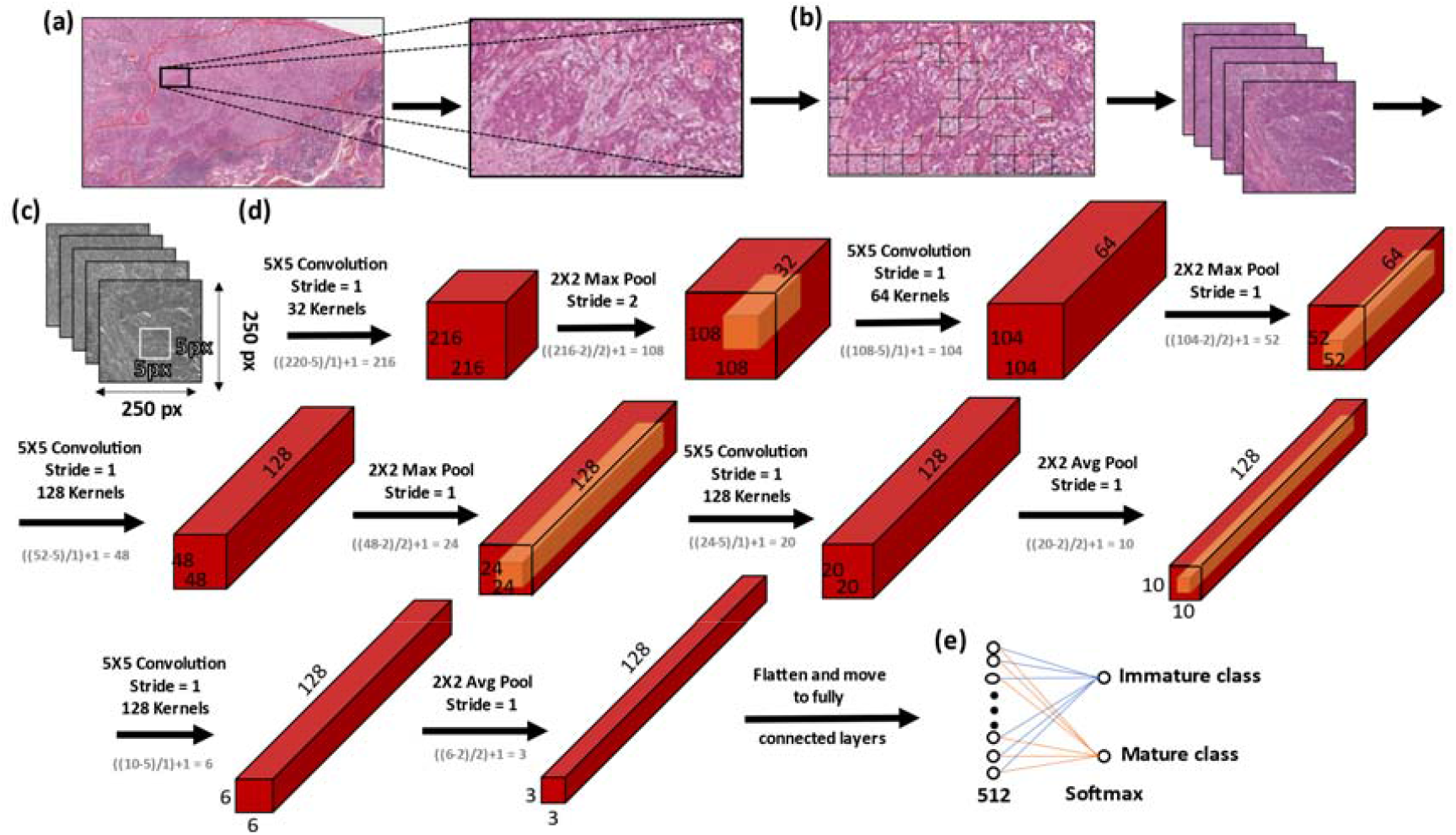
Workflow and architecture for the in-house convolutional neural network (CNN) used in the study. Regions of interest (ROI) are annotated and extracted from the tumour body (a). Image tiles of size 250×250 pixels were extracted from within stroma annotations within each ROI (b). Tiles were converted to greyscale to conserve memory, and fed into a CNN consisting of five convolutional layers interlinked with five pooling layers (c). A graphical representation of how these tiles are then processed within the five convolutional layers interlinked with five pooling layers of the CNN used in this study (d);the output of which is flattened before being fed into two fully connected layers wherein stromal maturity is predicted using the softmax function in the final layer (e). *(Avg = Average, Max = Maximum) Equations in grey show how feature map dimensions were calculated*.

### 2.7 Statistical analysis

The pathologist stromal maturity scores were used as the ground truth for development of the CNN. Model evaluation was conducted against the ground truth (pathologist scores) for the best-saved weights and bias in each of the image data sets at an individual tile, ROI and patient level. Confusion matrices were calculated to help determine the model’s precision, recall and F1-scores. Receiver-Operator Characteristic (ROC) curves were generated for assessment of the area under the curve (AUC) using the Scikit-learn library [28] in Python 3.8 at a tile and ROI level. Due to the heterogenous nature of some of the patients and methods of aggregation to predict outcome, ROC curves were not generated at this level.

Comparability between the best CNN model and the manual evaluation method was also assessed. Sensitivity, specificity, accuracy and their 95% confidence intervals were also calculated in the two additional independent manual stromal maturity classifications. For the purpose of this analysis, the model was considered a fourth evaluator. Inter-evaluator concordance was conducted using Fleiss’ Kappa. All bio-statistical analyses were performed using R v3.6.1[37].

## 3. Results

### 3.1 Patient Images

Classification of stromal maturity in digitally annotated ROI’s was conducted on H&E stained slides for 197 patients with OPSCC. From these patients, 636 ROIs were annotated and evaluated manually. In total, 9.91% (63/636) ROIs had insufficient stroma to produce tiles, resulting in 4.06% (8/197) patients being excluded from further analysis in the study. Of the remaining patients, 33.86% (64/189) were found to have immature stroma in all ROIs assessed and 45.50% (86/189) patients were found to have mature stroma present in all ROIs assessed. Classification of stromal maturity across ROIs was heterogeneous in 20.64% (39/189) of patients assessed. There were 29 heterogenous patients in the training group, 4 in the test group and 6 in the independent validation group (Figure 8).

**Figure 8.**
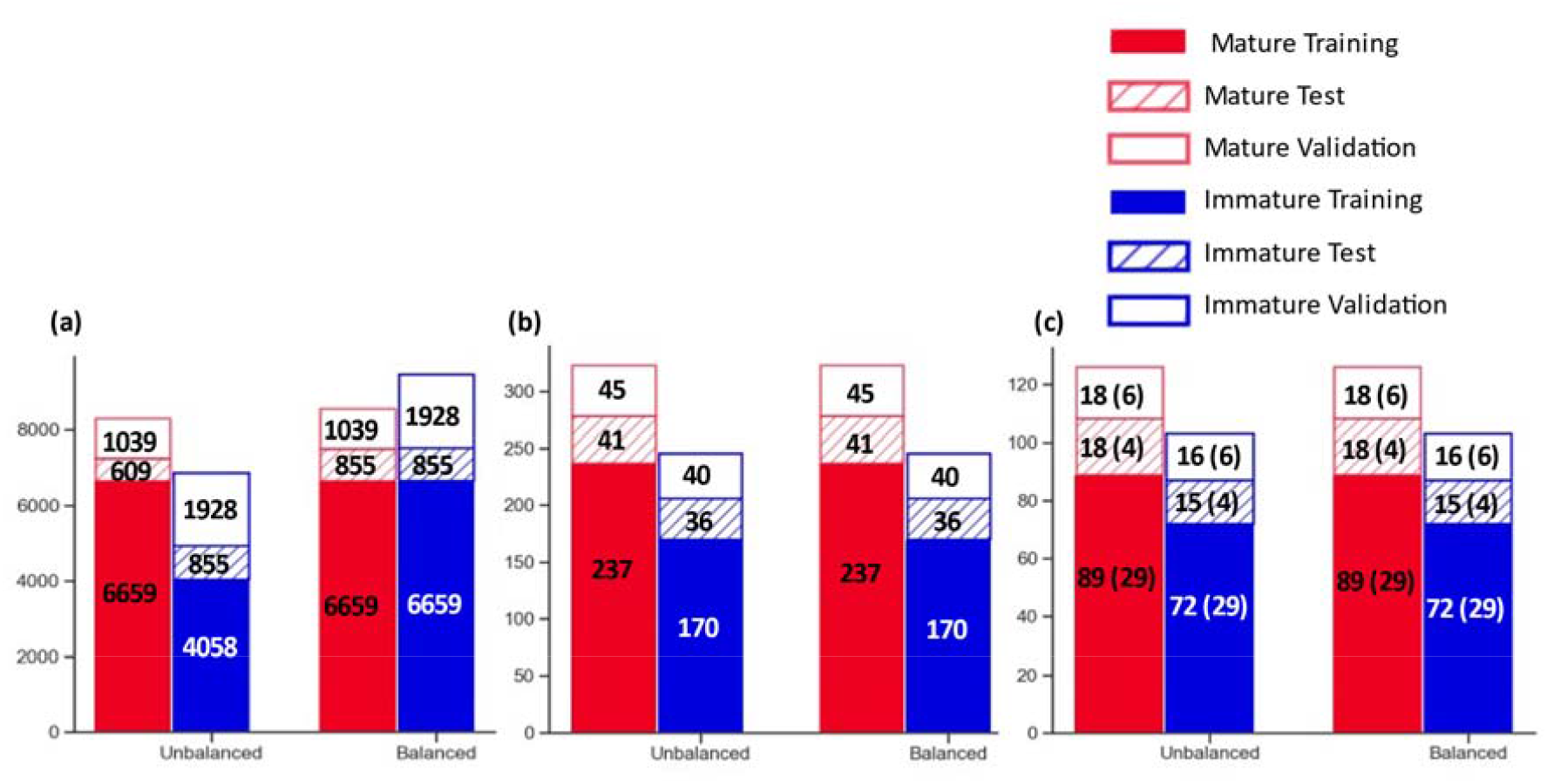
Histograms Showing breakdown of the image dataset for image tiles (A), ROIs (B) and at patient level (C) before and after dataset balancing. Datasets were only balanced at a tile level. NB The patient counts treat stromal heterogenous cases as both a mature an immature patient in these figures. The number of heterogenous patients are denoted in the parenthesis in (c)

### 3.2 Image Set Times

The time taken to perform each of the adjustments outlined in Table 1 were recorded for each image set. HistoClean balanced the baseline training data in 5.79 seconds with a difference of 2601 images translating to a rate of 452.35 images per second. Normalisation of all 15148 images in unbalanced training data took 66.85 seconds equating to a rate of 226.60 images per second. Embossing the unbalanced data took 30.45 seconds, a rate of 497.47 images per second.

The use of multithreading allowed for the processing of the images in a rapid timeframe. As mentioned previously, the number of threads used scales with the CPU cores, allowing the user to carry out other tasks while HistoClean produces the new images.

### 3.3 Evaluation of image data sets in robust CNN development

The CNN was trained eight separate times from scratch using the eight separate image sets summarised in Table 1. Use of image pre-processing techniques were found to consistently improve upon model performance when compared to the baseline “unbalanced” dataset across all levels of prediction assessed; from probability of individual image tiles to aggregation of probability at the patient level (Table 2). Image pre-processing conducted in the Balanced Embossed set provided the best overall accuracy at a tile, ROI and Patient level (0.774, 0.835 and 0.857 respectively) as well as a superior f1-score (0.820, 0.844 and 0.846 respectively). From these results, the balanced embossed set was determined to be the best preforming image set overall. In addition, the Balanced Embossed image set provided the best area under curve (AUC) scores (0.839 and 0.963 at a tile and patch level; Figure 9).

**Table 2.** Breakdown of model evaluation for each image set. Highlighted in bold are the best results for each category.

| Image Set | Level | True Mature (TN) | False Immature (FP) | True Immature (TP) | False Mature (FN) | Precision | Recall | F1 Score | Total Correct | Total Incorrect | ROC AUC | Overall Accuracy |
| --- | --- | --- | --- | --- | --- | --- | --- | --- | --- | --- | --- | --- |
| Unbalanced | Tile | 961 | 78 | 746 | 1182 | **0.905** | 0.387 | 0.542 | 1707 | 1260 | 0.791 | 0.575 |
|  | ROI | 41 | 4 | 12 | 28 | 0.750 | 0.300 | 0.429 | 53 | 32 | 0.636 | 0.624 |
|  | Patient | 14 | 1 | 3 | 10 | 0.750 | 0.231 | 0.353 | 17 | 11 | NA | 0.607 |
| Unbalanced Embossed | Tile | 733 | 306 | 1281 | 647 | 0.807 | 0.664 | 0.729 | 2014 | 953 | 0.757 | 0.679 |
|  | ROI | 29 | 16 | 34 | 6 | 0.680 | 0.850 | 0.756 | 63 | 22 | 0.882 | 0.741 |
|  | Patient | 10 | 5 | 10 | 3 | 0.667 | 0.769 | 0.714 | 20 | 8 | NA | 0.714 |
| Unbalanced Normalised | Tile | 817 | 222 | 1314 | 614 | 0.855 | 0.682 | 0.759 | 2131 | 836 | 0.811 | 0.718 |
|  | ROI | 34 | 11 | 35 | 5 | 0.761 | 0.875 | 0.814 | 69 | 16 | 0.884 | 0.812 |
|  | Patient | 11 | 4 | 11 | 2 | 0.786 | 0.769 | 0.777 | 20 | 8 | NA | 0.714 |
| Unbalanced Normalised Embossed | Tile | 772 | 267 | 1316 | 612 | 0.831 | 0.683 | 0.750 | 2088 | 879 | 0.782 | 0.704 |
|  | ROI | 33 | 12 | 37 | 3 | 0.755 | 0.925 | 0.831 | 70 | 15 | 0.886 | 0.824 |
|  | Patient | 12 | 3 | 11 | 2 | 0.786 | 0.846 | 0.815 | 23 | 5 | NA | 0.821 |
| Balanced | Tile | 847 | 192 | 1216 | 712 | 0.864 | 0.631 | 0.729 | 2063 | 904 | 0.811 | 0.695 |
|  | ROI | 37 | 8 | 30 | 10 | **0.789** | 0.750 | 0.769 | 67 | 18 | 0.86 | 0.788 |
|  | Patient | 14 | 1 | 9 | 4 | **0.900** | 0.692 | 0.783 | 24 | 5 | NA | 0.828 |
| Balanced Embossed | Tile | 764 | 275 | 1532 | 396 | 0.848 | 0.795 | **0.820** | 2296 | 671 | **0.839** | **0.774** |
|  | ROI | 33 | 12 | 38 | 2 | 0.760 | 0.950 | **0.844** | 71 | 14 | **0.963** | **0.835** |
|  | Patient | 13 | 2 | 11 | 2 | 0.846 | 0.846 | **0.846** | 24 | 4 | NA | **0.857** |
| Balanced Normalised | Tile | 742 | 297 | 1422 | 506 | 0.827 | 0.738 | 0.780 | 2164 | 803 | 0.798 | 0.729 |
|  | ROI | 32 | 13 | 36 | 4 | 0.735 | 0.900 | 0.809 | 68 | 17 | 0.897 | 0.800 |
|  | Patient | 11 | 4 | 11 | 2 | 0.733 | 0.846 | 0.786 | 22 | 6 | NA | 0.786 |
| Balanced Normalised Embossed | Tile | 623 | 416 | 1590 | 338 | 0.793 | **0.825** | 0.808 | 2213 | 754 | 0.788 | 0.746 |
|  | ROI | 27 | 18 | 39 | 1 | 0.684 | **0.975** | 0.804 | 66 | 19 | 0.932 | 0.776 |
|  | Patient | 9 | 6 | 13 | 0 | 0.684 | **1.000** | 0.813 | 22 | 6 | NA | 0.786 |

**Figure 9.**
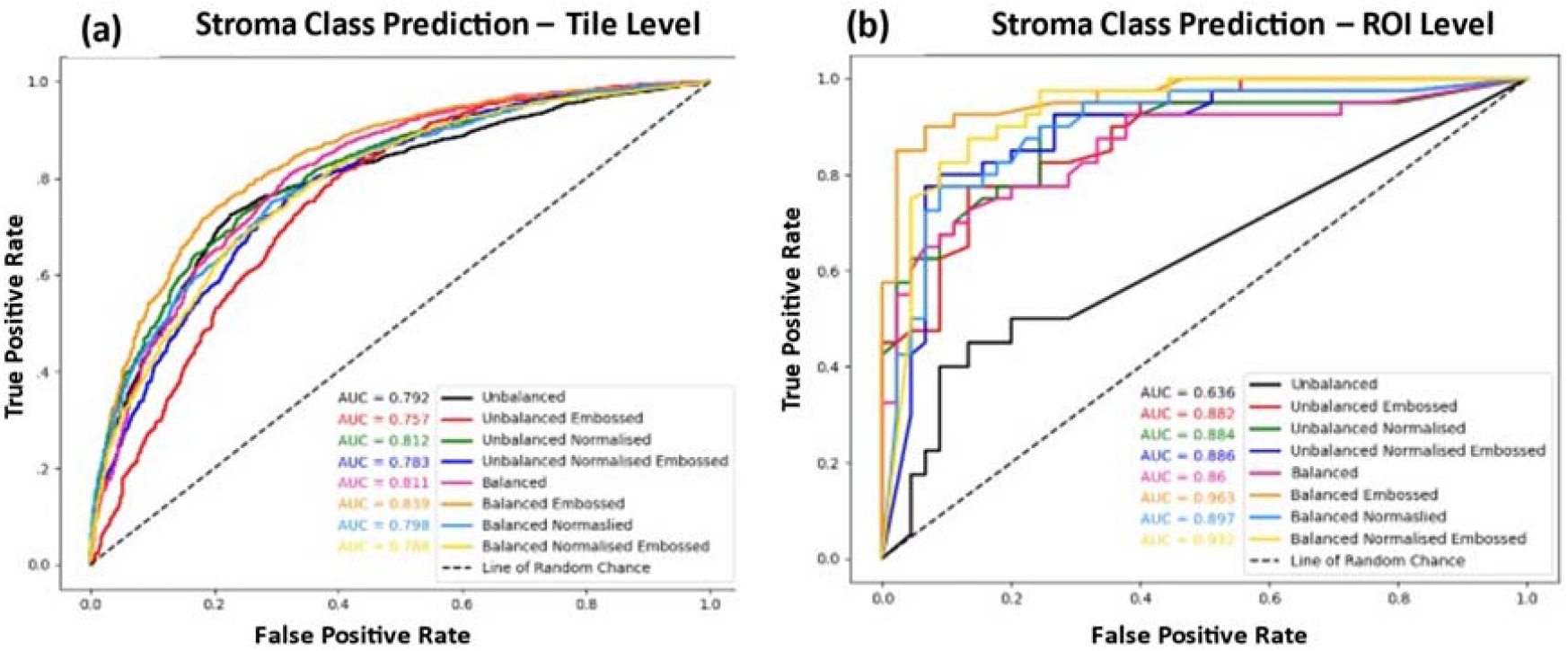
ROC Curve comparison of the different image datasets evaluated for CNN model accuracy within the image tiles (a) and ROIs (b). A combination of embossing and balancing the image sets provided the best overall area under curve (AUC) at a tile and ROI level.

The ability to predict stromal maturity using the CNN trained on the balanced embossed images was developed using the ground truth for stromal maturity in that ROI as provided by a single pathologist. Therefore, the sensitivity and specificity of manual classification of stromal maturity by two independent assessors to predict the pathologist scores was conducted and compared to results from balanced embossed image trained CNN in order to determine how reproducible the original pathologist scores were. Both independent manual assessors and the balanced embossed image set trained CNN demonstrated comparable sensitivity (100%; 95% CI, 77%–100%, for Assessor 1; 93%; 95% CI, 68%–100%, for Assessor 2 and 80%; 95% CI, 52%–96%, for the CNN) and specificity (86%; 95% CI, 57%– 98%, for Assessor 1; 100%; 95% CI, 75%–100%, for Assessor 2 and 85%; 95% CI, 55%– 98%, for the CNN) when classifying patients with having immature stroma based on the original pathologist scores. Moreover, the Fleiss’ Kappa score demonstrated good concordance between all three manual assessors and the CNN(κ = 0.785, p<0.0001). A review of misclassification by the balanced embossed image set trained CNN found misclassification occurred most often when a small number of tiles were available for stromal classification in that patient (Figure 10a). Misclassification by this model was found at a tile level whenever the image augmentation enhanced the presence of whitespace in immature stroma tiles resulting in misclassification of mature stroma in the embossed image (Figure 10b). In one patient, no tiles were able to be extracted from 3 of the 5 ROIs, resulting in an inversion of stromal maturity prediction that was subsequently incorrect.

**Figure 10.**
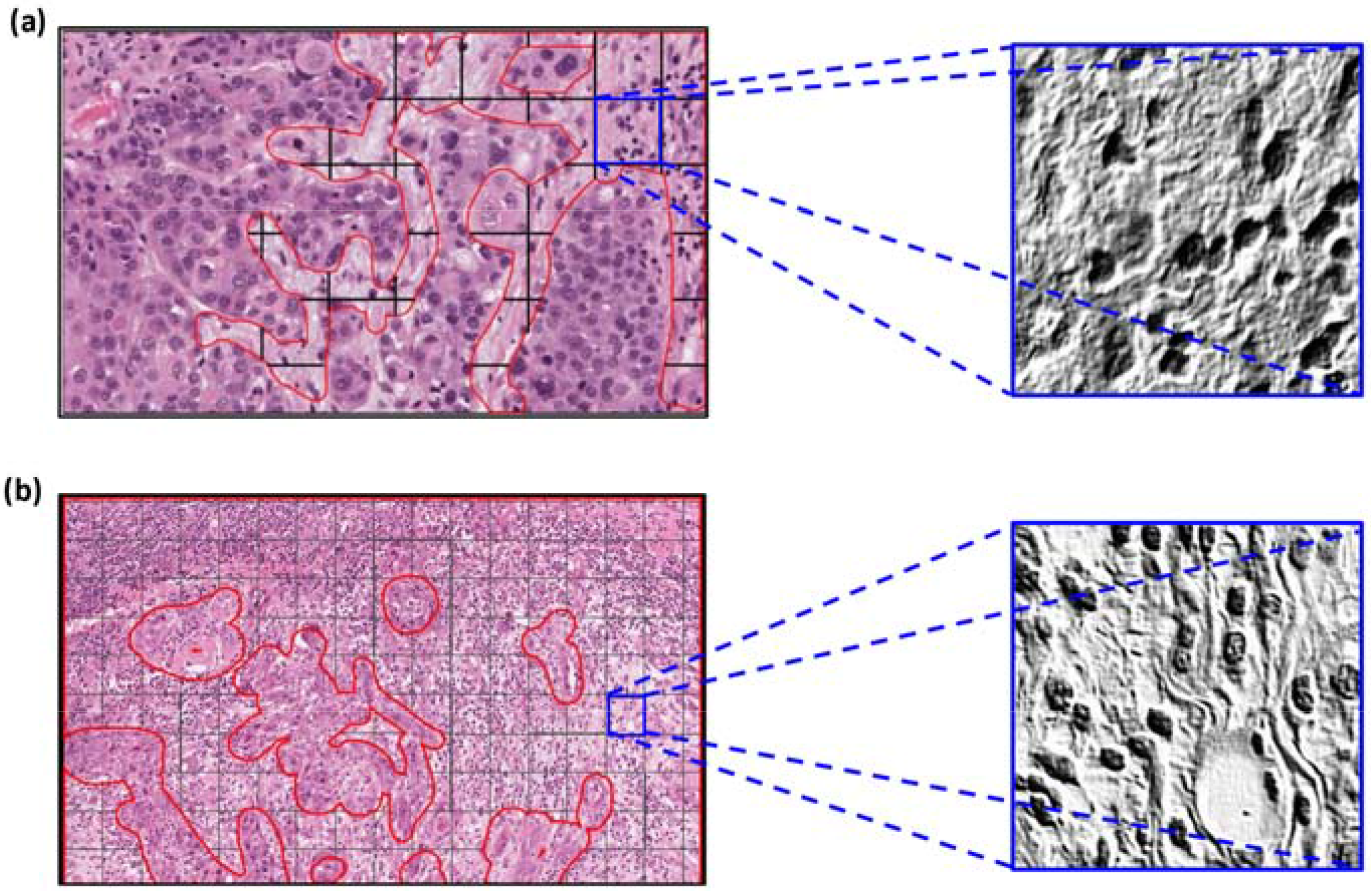
Representative examples of misclassified DS6 CNN stromal maturity prediction. Some patients in the cohort had limited stroma present, meaning very few tiles representative of overall patient’s stromal maturity could be extracted resulting in misclassification at a stromal independent patient level (a). Whilst at the tile level, image augmentation using the emboss technique was found to enhance linear structures surrounding oedema resulting in the embossed image possessing features associated with mature stroma resulting in misclassification of the tile (b).

## 4. Discussion

As technology advances, so too does the demand for computational, high-throughput, cost-effective diagnostic tools for use in clinical medicine. This is particularly true in the field of clinical pathology that traditionally has utilised fewer technological aids in spite of a depleting workforce (36, 37). Digital pathology, involves the acquisition and review of ultra-high-resolution whole slide images using a computer monitor in place of a microscope [40]. Digitisation of histological slides benefits from remote access for diagnostic reporting, providing a quick and easy means of recourse for diagnoses of complex pathology though ease of sharing virtual slides to consultant histopathologists with sub-specialist interest (38, 39). In addition, slide digitisation permits the use of digital image analysis tools to quantify histological features objectively using AI, as seen in radiomics [42]. At present, use of digital image analysis algorithms by consultant histopathologists is limited due to lack of modernisation in clinical pathology within the National Health Service, UK [41]. However, many consultant histopathologists recognise the benefit digital image analysis methodology could provide in streamlining the decision making process [43].

In contrast to other medical and non-medical disciplines that have implemented AI-assisted DIA, there is a scarcity of appropriate pathological images for developing deep learning models in clinical pathology [44]. This is in part due to the relatively recent move towards digitisation of pathology services, but more often due to lack of pathological material regarding the question of interest. Histological images are data rich and demonstrate significant heterogeneity across and within disease pathologies [45]. Therefore, the number of images required for effective deep learning is that of many orders of magnitude greater than that those required when developing models using more classical machine learning methods. Depending on the model being developed, this may require image datasets to be sourced at a global scale. Consequently, this introduces image variability and potential bias into CNN learning through differences in laboratory practice, scanning procedures or age of the sample being scanned [46]. This can have a pronounced effect on model learning and validation, particularly in small cohort studies, as each histological image possess unique image colour, contrasts and brightness profiles. The inter-laboratory variation limits the efficacy of developed models from small cohort students to be used in practice. CNNs have already shown promise in several cancer types and in several different use cases. One study by Khosravi et al. evaluated both in-house and the current top pretrained models’ efficacy across numerous cancer types and in several different tasks [47]. Many of these models achieved >90% accuracy in the categories of tumour detection, biomarker detection and tumour subtyping in bladder, breast and lung cancers. Another study demonstrated the use of several pretrained neural networks to identify different growth patterns in lung adenocarcinoma, achieving accuracies up to 85% [6].

In this study, we demonstrate the power of image pre-processing and augmentation and present a novel open-source GUI called HistoClean. Using a relatively simple CNN architecture, we clearly establish how use of image pre-processing techniques improves upon model generalisability for prediction of stromal maturity in an independent validation dataset. Further, we show that the best developed model, the balanced embossed model, had similar concordance, sensitivity and specificity to two further independent assessors of stromal maturity by manual review. However, we also show that poor choice of image pre-processing and augmentation techniques can introduce bias and noise. The use of image augmentation for dataset balancing helped to increase the small number of immature samples present for model development whilst image pre-processing through embossing helped to accentuate the features of interest we wanted the model to train with. Therefore, to ensure successful model development, consideration of which techniques to implement should reflect the specific research question being asked.

When trying to improve the accuracy of a CNN, often developmental time is spent refining the neural network and the network’s hyperparameters. However, it is arguably just as, if not more important to focus on the quality of the images used in training the network; a sentiment captured by the expression “rubbish in = rubbish out”. This study illustrates how crucial it is to balance the number of input images across the classes to prevent model overfitting. This initial step significantly improved both overall accuracy and AUC at the tile, patch and ROI level. The strength of this action is also clearly demonstrated by the change in false mature and false immature rates when comparing the balanced dataset to the unbalanced dataset. This is evidenced in the increases in f1-value at tile ROI and patient level (0.187, 0.340 and 0.443 respectively, Table 2). In parallel to this, embossing alone also demonstrated increases in accuracy and AUC across all levels, as well as lessening the effect of a mature dominant training set (Table 2). A synergistic improvement occurred when the dataset was both balanced and embossed, achieving an accuracy of 0.774 at a tile level. These improvements are in line with several other studies that use different augmentation techniques [48–50]. Importantly, HistoClean allowed the bio-image analyst to review the output of the image processing steps being applied within the software before proceeding to model development, providing opportunity for discussion of how particular image augmentations may enhance qualitative features the pathologist used to define stromal maturity in the image.

In this study, we also demonstrate that inappropriate augmentations can harm deep learning model development. This is evidenced by the reduction in accuracy between the Balanced Embossed and Balanced Normalised Embossed image sets, with a particular shift towards immature prediction as reflected in the increase in recall and decrease in precision at all levels. Upon examination of the patients in which this phenomenon had the greatest effect, it was clear that image normalisation, while correcting any colour imbalance, often created artefactual whitespace (Figure 11c). This was further highlighted by the embossing, (Figure 11d) causing the mature tiles to lose the dense parallel stromal fibres and adopt a more immature phenotype. This also raises the question of whether the improvements between the unbalanced and unbalanced normalised image sets are genuine or an artificial correction in the majority mature training data. It could be hypothesised that an immature skewed training set could suffer from further negative bias using this technique. Situations like this reinforce HistoClean as a useful tool for image pre-processing. A trained pathologist would be able to preview these changes and identify flaws in the pre-processing steps to avoid them. Furthermore, the traceability and data integrity provided by the application allows for easy comparison of the images.

**Figure 11.**
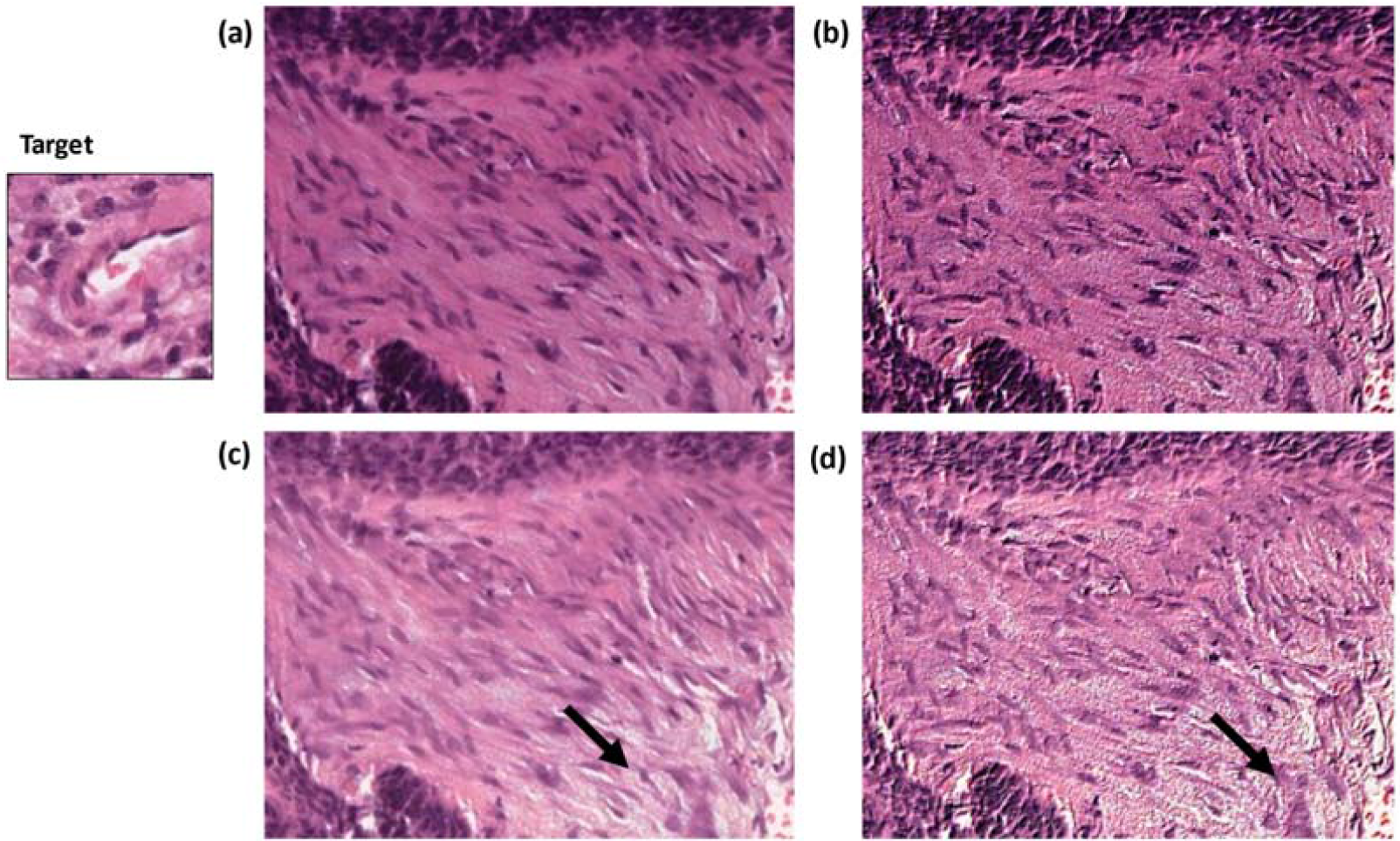
Example ground truth mature ROI. The original image (a) was embossed (b) and in the balanced embossed image set was predicted mature. Normalisation of the image created artefactual whitespace (c) which was then exacerbated by the embossing (d), flipping the prediction to an immature phenotype.

While the findings of this work give reason to be optimistic, there are still barriers to overcome before these tools are utilised in a clinical setting. With the common complaints of job losses and disconnect from the patient [51] aside, there can also be a lack of explainability and interpretability of the outcomes of neural networks; known as “Black Box” Deep learning [52]. This has led to a debate on how important it is to explain diagnostic outcome even if the accuracy is high [53]. However, this is comparable to the many commonly used drugs where we still lack a complete understanding of their mechanism of action [54]. There have been great efforts made to help uncover the logic behind image classification in deep learning models. These include the generation of saliency maps based on the generated gradients and loss [55], gradient-weighted class activation mapping [56], and minimal explainability maps ([57]). These techniques highlight areas of interest on the original images, providing some insight into which features are contributing to the classification. As techniques like this continue to improve, the concerns around the blind nature of deep learning should be alleviated.

## 5. Conclusions

This study confirms that use of image pre-processing and augmentation techniques available in HistoClean can advance the field of deep learning by facilitating arguably the most important step CNN-centric experiments; image set preparation. However, there is a lack of easy to use open-source software to facilitate this process. This study demonstrates the usefulness of HistoClean as an open-source software to implement image pre-processing techniques in image research, saving time and improving transparency and data integrity. HistoClean provides a rapid, robust and reproducible means of implementing these techniques in a way that can be used by experts, such as pathologists, to help identify which techniques could potentially be of use in their study.

## 6. Additional Information

## 6.1 Acknowledgments

The Northern Ireland OPSCC FFPE sections and linked clinicopathological data used in this research were received from the Northern Ireland Biobank, which has received funds from Health and Social Care Research and Development Division of the Public Health Agency in Northern Ireland and the Friends of the Cancer Centre. The Precision Medicine Centre of Excellence has received funding from Invest Northern Ireland, Cancer Research UK, Health and Social Care Research and Development Division of the Public Health Agency in Northern Ireland, Seán Crummey Memorial Fund, and Tom Simms Memorial Fund.

## Abbreviations

AI: Artificial Intelligence
DIA: Digital Image Analysis
GUI: Graphical User Interface
ROC: Receiver-Operator Characteristic
AUC: Area Under Curve

## References

[1] Bankhead P, Loughrey MB, Fernández JA, Dombrowski Y, McArt DG, Dunne PD, et al. QuPath: Open source software for digital pathology image analysis. Sci Rep 2017;7:1–7. https://doi.org/10.1038/s41598-017-17204-5.

[2] Satyanarayanan M, Goode A, Gilbert B, Harkes J, Jukic D. OpenSlide: A vendor-neutral software foundation for digital pathology. J Pathol Inform 2013;4:27. https://doi.org/10.4103/2153-3539.119005.

[3] Rueden CT, Schindelin J, Hiner MC, Dezonia BE, Walter AE, Arena ET, et al. ImageJ2□: ImageJ for the next generation of scientific image data 2017:1–26. https://doi.org/10.1186/s12859-017-1934-z.

[4] Salto-Tellez M, Maxwell P, Hamilton P. Artificial intelligence-the third revolution in pathology. Histopathology 2019;74:372–6. https://doi.org/10.1111/his.13760.

[5] Kim KG. Book Review: Deep Learning. Healthc Inform Res 2016;22:351. https://doi.org/10.4258/hir.2016.22.4.351.

[6] Gertych A, Swiderska-chadaj Z, Ma Z, Ing N, Markiewicz T, Cierniak S, et al. Convolutional neural networks can accurately distinguish four histologic growth patterns of lung adenocarcinoma in digital slides. Sci Rep 2019:1–12. https://doi.org/10.1038/s41598-018-37638-9.

[7] Dosovitskiy A, Fischer P, Springenberg JT, Riedmiller M, Brox T. Discriminative unsupervised feature learning with exemplar convolutional neural networks. IEEE Trans Pattern Anal Mach Intell 2016;38:1734–47. https://doi.org/10.1109/TPAMI.2015.2496141.

[8] Zhao R, Luk W, Niu X, Shi H, Wang H. Hardware Acceleration for Machine Learning 2017:2–7. https://doi.org/10.1109/ISVLSI.2017.127.

[9] von Chamier L, Laine RF, Jukkala J, Spahn C, Krentzel D, Nehme E, et al. Democratising deep learning for microscopy with ZeroCostDL4Mic. Nat Commun 2021;12:2276. https://doi.org/10.1038/s41467-021-22518-0.

[10] Gómez-de-Mariscal E, García-López-de-Haro C, Donati L, Unser M, Munoz-Barrutia A, Sage D. DEEPIMAGEJ: A USER-FRIENDLY PLUGIN TO RUN DEEP LEARNING MODELS IN IMAGEJ. BioRxiv 2019:1–13. https://doi.org/ https://doi.org/10.1101/799270.

[11] Janowczyk A, Zuo R, Gilmore H, Feldman M, Madabhushi A. HistoQC: An Open-Source Quality Control Tool for Digital Pathology Slides. JCO Clin Cancer Informatics 2019:1–7. https://doi.org/10.1200/cci.18.00157.

[12] Kourou K, Exarchos TP, Exarchos KP, Karamouzis M V., Fotiadis DI. Machine learning applications in cancer prognosis and prediction. Comput Struct Biotechnol J 2015;13:8–17. https://doi.org/10.1016/j.csbj.2014.11.005.

[13] Balkenhol M, Karssemeijer N, Litjens GJS, van der Laak J, Ciompi F, Tellez D. H&E stain augmentation improves generalization of convolutional networks for histopathological mitosis detection 2018:34. https://doi.org/10.1117/12.2293048.

[14] Zhu C, Song F, Wang Y, Dong H, Guo Y, Liu J. Breast cancer histopathology image classification through assembling multiple compact CNNs. BMC Med Inform Decis Mak 2019;19:1–17. https://doi.org/10.1186/s12911-019-0913-x.

[15] Kassani SH, Kassani PH, Wesolowski MJ, Schneider KA, Deters R. Classification of histopathological biopsy images using ensemble of deep learning networks. CASCON 2019 Proc - Conf Cent Adv Stud Collab Res - Proc 29th Annu Int Conf Comput Sci Softw Eng 2020:92–9.

[16] Komura D, Ishikawa S. Machine Learning Methods for Histopathological Image Analysis. Comput Struct Biotechnol J 2018;16:34–42. https://doi.org/10.1016/j.csbj.2018.01.001.

[17] Hensman P, Masko D. The Impact of Imbalanced Training Data for Convolutional Neural Networks. PhD 2015. https://www.diva-portal.org/smash/get/diva2:811111/FULLTEXT01.pdf%C3%AF%C2%BC%E2%80%B0 (accessed June 7, 2021)

[18] Khan AM, Rajpoot N, Treanor D, Magee D. A nonlinear mapping approach to stain normalization in digital histopathology images using image-specific color deconvolution. IEEE Trans Biomed Eng 2014;61:1729–38. https://doi.org/10.1109/TBME.2014.2303294.

[19] Mikołajczyk A, Grochowski M. Data augmentation for improving deep learning in image classification problem. 2018 Int Interdiscip PhD Work IIPhDW 2018 2018:117– 22. https://doi.org/10.1109/IIPHDW.2018.8388338.

[20] Abadi M, Agarwal A, Barham P, Brevdo E, Chen Z, Citro C, et al. TensorFlow: Large-Scale Machine Learning on Heterogeneous Distributed Systems 2016. https://doi.org/ https://doi.org/10.1101/2020.03.20.000133.

[21] Tuininga A. cx-Freeze 2020. https://cx-freeze.readthedocs.io/en/latest/index.html. (accessed June 7, 2021)

[22] Hashemi M. Enlarging smaller images before inputting into convolutional neural network□: zero □ padding vs. interpolation. J Big Data 2019. https://doi.org/10.1186/s40537-019-0263-7.

[23] Weiyuan W, Verma D, Yang W. Patchify Github Repository. GitHub n.d. https://pypi.org/project/patchify/.(accessed June 7, 2021)

[24] Clark DP. A Deep Learning Convolutional Neural Network Can Recognize Common Patterns of Injury in Gastric 2020;144. https://doi.org/10.5858/arpa.2019-0004-OA.

[25] Goh TY, Basah SN, Yazid H, Juhairi M, Safar A. Performance analysis of image thresholding□: Otsu technique. Measurement 2018;114:298–307. https://doi.org/10.1016/j.measurement.2017.09.052.

[26] Bradski G. The OpenCV Library. Dr Dobb’s J Softw Tools 2000.

[27] Mikolajczyk A, Grochowski M. Data augmentation for improving deep learning in image classification problem. 2018 Int. Interdiscip. PhD Work., IEEE; 2018, p. 117– 22. https://doi.org/10.1109/IIPHDW.2018.8388338.

[28] Walt V Der, Sch JL, Nunez-iglesias J. scikit-image□: image processing in Python 2014:1–18. https://doi.org/10.7717/peerj.453.

[29] Gonzalez RC, Woods RE. Digital Image Processing. 2018.

[30] Litjens G, Kooi T, Bejnordi BE, Setio AAA, Ciompi F, Ghafoorian M, et al. A survey on deep learning in medical image analysis. Med Image Anal 2017;42:60–88. https://doi.org/10.1016/j.media.2017.07.005.

[31] Sudeep KS, Pal KK. Preprocessing for image classification by convolutional neural networks. 2016 IEEE Int Conf Recent Trends Electron Inf Commun Technol RTEICT 2016 - Proc 2017:1778–81. https://doi.org/10.1109/RTEICT.2016.7808140.

[32] Jung AB, Crall J, Wada K, Tanaka S, Graving J, Reinders C, et al. imgaug. Online 2020. https://github.com/aleju/imgaug (accessed November 25, 2020).

[33] Craig SG, Anderson LA, Moran M, Graham L, Currie K, Rooney K, et al. Comparison of molecular assays for HPV testing in oropharyngeal squamous cell carcinomas: A population-based study in Northern Ireland. Cancer Epidemiol Biomarkers Prev 2020;29:31–8. https://doi.org/10.1158/1055-9965.EPI-19-0538.

[34] Ueno H. Histological categorisation of fibrotic cancer stroma in advanced rectal cancer. Gut 2004;53:581–6. https://doi.org/10.1136/gut.2003.028365.

[35] Kemi NA, Eskuri M, Pohjanen VM, Karttunen TJ, Kauppila JH. Histological assessment of stromal maturity as a prognostic factor in surgically treated gastric adenocarcinoma. Histopathology 2019;75:882–9. https://doi.org/10.1111/his.13934.

[36] Paszke A, Gross S, Massa F, Lerer A, Bradbury J, Chanan G, et al. PyTorch: An Imperative Style, High-Performance Deep Learning Library. ArXiv 2019.

[37] Core Team R. R: A Language and Environment for Statistical Computing 2021. https://www.r-project.org/.

[38] Metter DM, Colgan TJ, Leung ST, Timmons CF, Park JY. Trends in the US and Canadian Pathologist Workforces From 2007 to 2017. JAMA Netw Open 2019;2:e194337. https://doi.org/10.1001/jamanetworkopen.2019.4337.

[39] Bainbridge S, Cake R, Mike M, Furness P, Gordon B. Testing Times To Come□? An Evaluation of Pathology Capacity Across the Uk. Cancer Res UK 2016.

[40] Jahn SW, Plass M, Moinfar F. Digital Pathology: Advantages, Limitations and Emerging Perspectives. J Clin Med 2020;9:3697. https://doi.org/10.3390/jcm9113697.

[41] Griffin J, Treanor D. Digital pathology in clinical use: Where are we now and what is holding us back? Histopathology 2017;70:134–45. https://doi.org/10.1111/his.12993.

[42] van Timmeren JE, Cester D, Tanadini-Lang S, Alkadhi H, Baessler B. Radiomics in medical imaging—”how-to” guide and critical reflection. Insights Imaging 2020;11. https://doi.org/10.1186/s13244-020-00887-2.

[43] Williams BJ, Bottoms D, Treanor D. Future-proofing pathology: The case for clinical adoption of digital pathology. J Clin Pathol 2017;70:1010–8. https://doi.org/10.1136/jclinpath-2017-204644.

[44] Anwar SM, Majid M, Qayyum A, Awais M, Alnowami M, Khan MK. Medical Image Analysis using Convolutional Neural Networks: A Review. J Med Syst 2018;42:226. https://doi.org/10.1007/s10916-018-1088-1.

[45] Bosman FT. Tumor Heterogeneity□: Will It Change What Pathologists Do□? 2018:18–22. https://doi.org/10.1159/000469664.

[46] Roy S, kumar Jain A, Lal S, Kini J. A study about color normalization methods for histopathology images. Micron 2018;114:42–61. https://doi.org/10.1016/j.micron.2018.07.005.

[47] Khosravi P, Kazemi E, Imielinski M. EBioMedicine Deep Convolutional Neural Networks Enable Discrimination of Heterogeneous Digital Pathology Images. EBioMedicine 2018;27:317–28. https://doi.org/10.1016/j.ebiom.2017.12.026.

[48] Salehinejad H, Colak E, Dowdell T, Barfett J, Valaee S. Synthesizing Chest X-Ray Pathology for Training Deep Convolutional Neural Networks. IEEE Trans Med Imaging 2019;38:1197–206. https://doi.org/10.1109/TMI.2018.2881415.

[49] Sajjad M, Khan S, Muhammad K, Wu W, Ullah A, Baik SW. Multi-grade brain tumor classification using deep CNN with extensive data augmentation. J Comput Sci 2019;30:174–82. https://doi.org/10.1016/j.jocs.2018.12.003.

[50] Tellez D, Litjens G, Bándi P, Bulten W, Bokhorst JM, Ciompi F, et al. Quantifying the effects of data augmentation and stain color normalization in convolutional neural networks for computational pathology. Med Image Anal 2019;58. https://doi.org/10.1016/j.media.2019.101544.

[51] Amisha, Malik P, Pathania M, Rathaur V. Overview of artificial intelligence in medicine. J Fam Med Prim Care 2019;8:2328. https://doi.org/10.4103/jfmpc.jfmpc_440_19.

[52] Adadi A, Berrada M. Peeking Inside the Black-Box: A Survey on Explainable Artificial Intelligence (XAI). IEEE Access 2018;6:52138–60. https://doi.org/10.1109/ACCESS.2018.2870052.

[53] London AJ. Artificial Intelligence and Black-Box Medical Decisions: Accuracy versus Explainability. Hastings Cent Rep 2019;49:15–21. https://doi.org/10.1002/hast.973.

[54] Gregori-puigjané E, Setola V, Hert J, Crews BA, Irwin JJ, Lounkine E. Identifying mechanism-of-action targets for drugs and probes 2012;109. https://doi.org/10.1073/pnas.1204524109.

[55] Simonyan K. Deep Inside Convolutional Networks□: Visualising Image Classification Models and Saliency Maps arXiv□: 1312. 6034v2 [cs. CV] 19 Apr 2014 2013:1–8.

[56] Selvaraju RR, Cogswell M, Das A, Vedantam R, Parikh D, Batra D. Grad-CAM: Visual Explanations from Deep Networks via Gradient-Based Localization. Int J Comput Vis 2020;128:336–59. https://doi.org/10.1007/s11263-019-01228-7.

[57] Sun Y, Chockler H, Huang X, Kroening D. Explaining Image Classifiers Using Statistical Fault Localization. In: Vedaldi A, Bischof H, Brox T, Frahm J-M, editors. Comput. Vis. -- ECCV 2020, Cham: Springer International Publishing; 2020, p. 391–406.

