## Supplemental Table 1 for "HistoClean: Open-source Software for Histological Image Pre-processing and Augmentation to Improve Development of Robust Convolutional Neural Networks"

### Slide 1
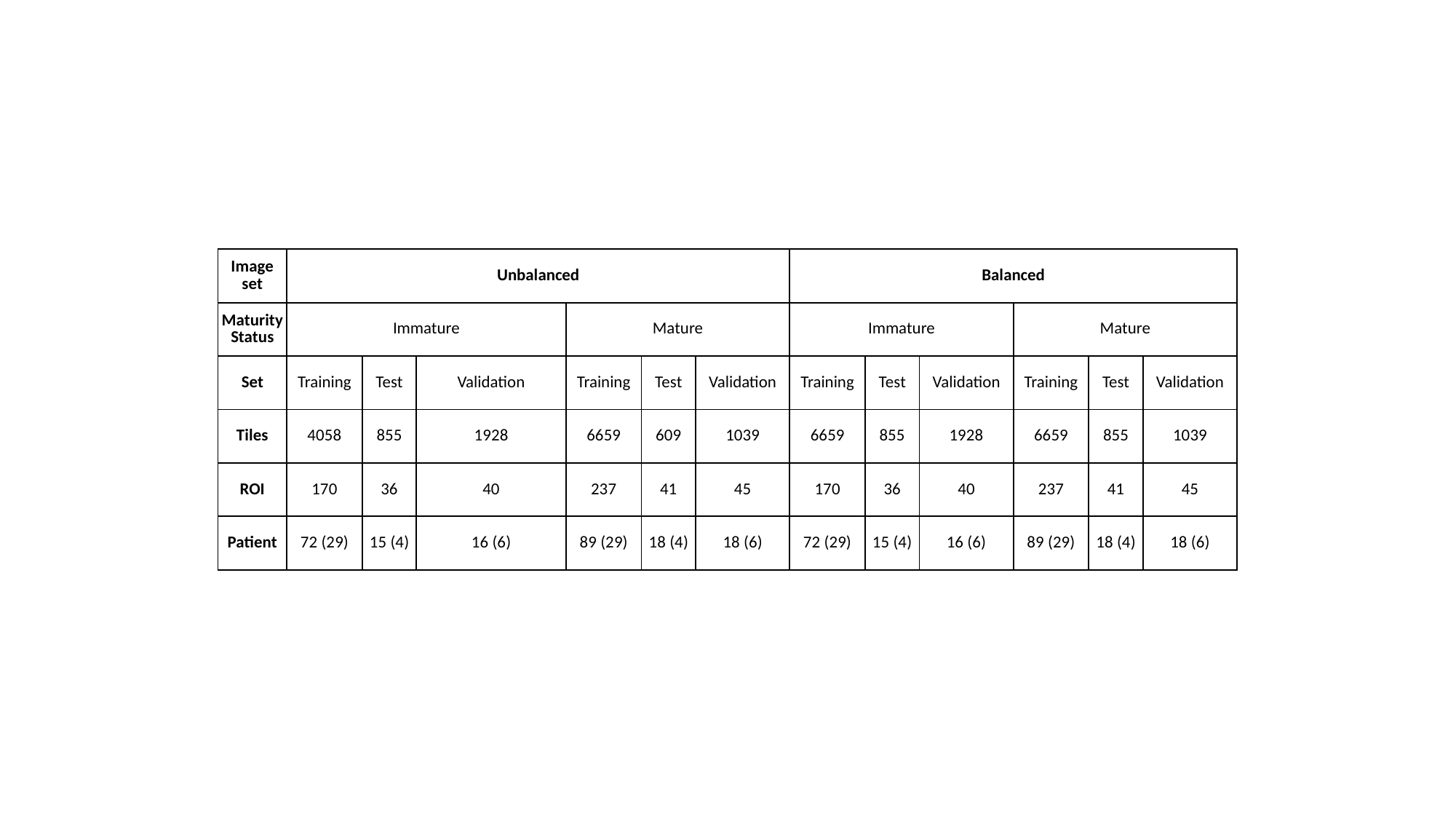

| Image set | Unbalanced | | | | | | Balanced | | | | | |
| --- | --- | --- | --- | --- | --- | --- | --- | --- | --- | --- | --- | --- |
| Maturity Status | Immature | | | Mature | | | Immature | | | Mature | | |
| Set | Training | Test | Validation | Training | Test | Validation | Training | Test | Validation | Training | Test | Validation |
| Tiles | 4058 | 855 | 1928 | 6659 | 609 | 1039 | 6659 | 855 | 1928 | 6659 | 855 | 1039 |
| ROI | 170 | 36 | 40 | 237 | 41 | 45 | 170 | 36 | 40 | 237 | 41 | 45 |
| Patient | 72 (29) | 15 (4) | 16 (6) | 89 (29) | 18 (4) | 18 (6) | 72 (29) | 15 (4) | 16 (6) | 89 (29) | 18 (4) | 18 (6) |
